# Germline testing data validate inferences of mutational status for variants detected from tumor-only sequencing

**DOI:** 10.1101/2021.04.14.439855

**Authors:** Nahed Jalloul, Israel Gomy, Samantha Stokes, Alexander Gusev, Bruce E. Johnson, Neal I. Lindeman, Laura Macconaill, Shridar Ganesan, Judy E. Garber, Hossein Khiabanian

**Author notes:** **Corresponding address:** Hossein Khiabanian, PhD, Associate Professor of Pathology in Medical Informatics Center for Systems and Computational Biology, Rutgers Cancer Institute of New Jersey, Rutgers University 195 Little Albany Street, New Brunswick, NJ, 08903-2681. These authors contributed equally to this work (Nahed Jalloul, Israel Gomy). These authors were co-leaders of this work (Judy E. Garber, Hossein Khiabanian).

## Abstract

**Background:** Pathogenic germline variants (PGV) in cancer susceptibility genes are usually identified in cancer patients through germline testing of DNA from blood or saliva: their detection can impact patient treatment options and potential risk reduction strategies for relatives. PGV can also be identified, in tumor sequencing assays, often performed without matched normal specimens. It is then critical to determine whether detected variants are somatic or germline. Here, we evaluate the clinical utility of computational inference of mutational status in tumor-only sequencing compared to germline testing results.

**Patients and Methods:** Tumor-only sequencing data from 1,608 patients were retrospectively analyzed to infer germline-versus-somatic status of variants using an information-theoretic, gene-independent approach. Loss of heterozygosity (LOH) was also determined. The predicted mutational models were compared to clinical germline testing results. Statistical measures were computed to evaluate performance.

**Results:** Tumor-only sequencing detected 3,988 variants across 70 cancer susceptibility genes for which germline testing data were available. Our analysis imputed germline-versus-somatic status for >75% of all detected variants, with a sensitivity of 65%, specificity of 88%, and overall accuracy of 86% for pathogenic variants. False omission rate was 3%, signifying minimal error in misclassifying true PGV. A higher portion of PGV in known hereditary tumor suppressors were found to be retained with LOH in the tumor specimens (72%) compared to variants of uncertain significance (58%).

**Conclusions:** Tumor-only sequencing provides sufficient power to distinguish germline and somatic variants and infer LOH. Although accurate detection of PGV from tumor-only data is possible, analyzing sequencing data in the context of specimens’ tumor cell content allows systematic exclusion of somatic variants, and suggests a balance between type 1 and 2 errors for identification of patients with candidate PGV for standard germline testing. Our approach, implemented in a user-friendly bioinformatics application, facilities objective analysis of tumor-only data in clinical settings.

**Highlights:**

- Most pathogenic germline variants in cancer predisposition genes can be identified by analyzing tumor-only sequencing data.
- Information-theoretic gene-independent analysis of common sequencing data accurately infers germline vs. somatic status.
- A reasonable statistical balance can be established between sensitivity and specificity demonstrating clinical utility.
- Pathogenic germline variants are more often detected with loss of heterozygosity vs. germline variants of uncertain significance.

## INTRODUCTION

Precision oncology relies on robust molecular analyses of patient samples and accurate interpretation of genomic sequencing and biomarker data to guide treatment strategies [1]. Technological advances in genomic sequencing have made tumor genomic profiling a routine process in the clinical evaluation and treatment planning of cancer patients [2]. The main objective of sequencing is to provide a detailed genomic characterization of the patient’s neoplasm, improve predictions on clinical outcome, and identify and potentially target oncogenic drivers to enable the development of an individualized treatment plan [3].

A small but important set of cancers arise in patients with pathogenic germline variants (PGV) that can both inform personal and familial cancer risks and guide treatment approaches [4]. Clinical germline testing has typically been limited to patients with personal and/or family history of tumors highly suggestive of specific predisposition syndromes. Germline DNA is analyzed for pathogenic alterations in one or more specific gene(s). However, germline testing is now expanding to a larger group of patients beyond those with a compelling family history [5]. Previously, effort was made to test individuals for PGV in only those genes most likely to confer risk consistent with the personal and family cancer history. However, cancer phenotypes may overlap among syndromes, and gene-sets may be under- or overrepresented in some panels. In addition, for a patient to be referred for clinical germline testing, certain features are often required by health insurance companies, which can restrict uptake. Because of complexities in determining the need for clinical germline testing, eligible patients are frequently overlooked and not tested [6]. A recent study showed that one in eight adult cancer patients who underwent universal germline testing, regardless of the extent to which they met established criteria, had a PGV in a susceptibility gene [7]. Almost half of these PGVs would not have been identified if testing criteria had been followed. Further, approximately one third of the PGV carriers had their therapies changed as a result.

Tumor sequencing for the identification of somatic alterations is becoming more widely carried out in patients with different cancers [8, 9]. Many commercial and academic tumor sequencing assays include a large set of cancer-related genes (>50) that can be mutated somatically and also confer cancer risk when mutated in the germline. To definitively identify somatic variants and potential germline variants in cancer cells, some laboratories analyze matched tumor and non-tumor specimens (e.g. blood, buccal mucosa, adjacent tissue) [8]. Multiple studies have shown that integration of tumor sequencing and matched normal genomic profiling can identify PGV in cancer predisposition genes in 15–18% of cancer patients, including those without high-risk family history or otherwise meeting clinical criteria for standard germline testing [10, 11]. These data suggest that current germline testing strategies may miss a significant number of mutation carriers in the population that are not identified by the patients’ and/or family history.

While concomitant tumor and germline sequencing analyses for all cancer patients may eventually become the standard of care in the future, an objective and reliable means of identifying patients for clinical germline testing confirmation is needed in the clinic today [12]. Current practice in interpreting tumor-only data for this purpose are gene-specific and are often based on variant allele frequency criteria that may need to be adjusted for different settings [13, 14]. To address these needs, we examined the performance of a gene-independent, information-theoretic pipeline aimed at accurately categorizing the variants identified by tumor-only assays as somatic or germline. Using commonly available sequencing data, we analyzed each variant in the context of specimen’s proportion of tumor cells and utilized high-depth sequencing to predict loss of heterozygosity status, which can potentially inform the functional effect of the mutation in both germline and somatic variants.

## METHODS

### Patient cohort and sample data

The cohort included a total of 1,608 patients with diverse malignancies who were consented to the PROFILE study [15] (protocols 11-104 and 17-000) at Dana-Farber Cancer Institute between January 2014 to December 2018 and had undergone somatic sequencing in the Center for Advanced Molecular Diagnostics at the Brigham and Women’s Hospital, and clinical germline testing. Genomic DNA was isolated from formalin-fixed paraffin embedded (FFPE) tissues containing at least 20% tumor nuclei and analyzed using the OncoPanel assay, which utilizes the Agilent SureSelect hybrid capture kit and Illumina HiSeq massive parallel sequencer according to standard pipelines as previously described [15]. The panel interrogates all exons and 191 introns in 447 genes to detect single nucleotide variants, indels, copy-number alterations, and structural variants. Germline testing and reporting were carried out by CLIA-certified commercial laboratories from blood samples collected clinically with consent, following the current American College of Medical Genetic (ACMG) guidelines [16].

Retrospective clinical, demographic data, and genomic data were accessed and de-identified through HIPAA-compliant IRB-approved chart review. Clinical information (age, sex, tumor type) and tumor-only sequencing data from the OncoPanel assay included altered genes, amino acid changes, cDNA changes, variant positions, reference and altered alleles, variant classifications, variant types, variant allele frequencies (VAF), copy-numbers, and sequencing depths, as well as histological estimates for proportion of tumor cells. Germline testing results for the corresponding patient samples included genes interrogated in the specific panel from one of five commercial testing laboratories (Ambry Genetics, Aliso Viejo, CA; Color Genomics, Burlingame, CA; GeneDX, Gaithersburg, MD; Invitae Corporation, San Francisco, CA; Myriad Genetics, Salt Lake City, UT). Variants of uncertain significance (VUS) were considered true germline. Variants classified as likely benign or benign are not routinely reported and were not included. VAF were obtained clinically by a genetic counselor for the *TP53* variants to aid in clinically distinguishing germline from acquired mosaicism or clonal hematopoiesis, both increasingly observed in cancer patients following exposure to cytotoxic chemotherapy and other risk factors [17]. Nomenclature variations between tumor sequencing and germline testing data were curated for 70 overlapping genes between the assays by comparing the reference transcript number, the position and type of the alteration in the specific genes.

### Tumor sequencing data analyses

The proportion of tumor cells (purity) and its confidence intervals were computationally estimated for all specimens using All-FIT (Allele-Frequency-based Imputation of Tumor Purity) [18]. To impute tumor cell content, All-FIT uses VAF of all SNV and indels, and sequencing depth and copy-number at their genomic position, which are commonly available in clinical tumor sequencing reports. Computational estimates were significantly correlated with specimen histological assessments of tumor purity (Pearson *r* = 0.31; p <0.001).

Next, both the histological purity and computational purity estimates were used to infer germline versus somatic mutational status and evaluate loss of heterozygosity for SNV and indels using LOHGIC (Loss of Heterozygosity Germline Inference Calculator) [19]. LOHGIC calculates weights for the likelihood of each somatic and germline mutational model (**Supplementary Figure 1**), taking into account the uncertainties in estimates of tumor purity and VAF measurements, which depend on sequencing depth. The most consistent model for a variant was selected based on the sum of the weights from all germline models (*W*_germ_) versus those from all somatic models (*W*_som_). Using criteria determined from simulations [19], when *W*_germ_ > 0.7 for a variant, it was inferred as germline; when *W*_som_ > 0.7 for a variant, it was inferred as somatic. If neither the *W*_germ_ nor *W*_som_ were greater than 0.7 for a variant, inference status was marked as ambiguous. Sum of the weights for germline or somatic LOH models larger than 0.5 was considered as evidence for the presence of LOH in the tumor.

Undetected focal copy-number alterations, inaccurate purity estimates, and low sequencing depths may result in ambiguous inferences. Specifically, large confidence intervals for VAF arising from low sequencing depths can produce cofounding results. For example, in a specimen with purity of 0.6 (assuming 5% inaccuracy), at sequencing depth of 1000, the confidence interval for an observed VAF of 0.5 is between 0.46 and 0.54. The largest weight for such a measurement would be 0.8 for a germline heterozygous model. However, for the same purity and observed VAF, sequencing at depth of 200 results in a larger confidence interval (0.42–0.58) and confounding weights across multiple models: *W* of 0.35 for somatic under LOH, *W* of 0.19 for somatic copy-neutral homozygous, and *W* of 0.45 for germline heterozygous, neither of which are sufficiently large for non-ambiguous inference.

Genomics Oncology Platform is a python GUI, freely available for the extraction of relevant information and the application of All-FIT and LOHGIC directly on variant calls. A snapshot of the application, showing variant data and status, with option to visualize mutational inferences represented by VAF vs. purity graphs is shown in **Supplementary Figure 1**. This application and individual algorithms are available at software.khiabanian-lab.org.

Pathogenicity of germline variants was assessed using curated open-access FDA-approved knowledge bases (ClinVar/ClinGen) and variants with conflicting interpretation of pathogenicity were manually curated using the ACMG guidelines. PGV included both pathogenic and likely pathogenic variants according to the ACMG classification [16]. Pathogenic status and mutational effect of somatic alterations were assessed using the oncoKB annotator, a precision oncology knowledge base [20], which provides the biological effect, prevalence and prognostic information, as well as treatment implications of alterations present in 682 cancer genes. We considered variants annotated as oncogenic, likely oncogenic, and predicted oncogenic as pathogenic.

### Statistical evaluation

Statistical measures computed for detected variants included: true positive rate or sensitivity, true negative rate or specificity, accuracy, positive predictive value or precision, false omission rate, and the F-score. These measures were used to evaluate the performance of the predicted mutational status using LOHGIC, inferred by both histological and computational purity estimates. However, in imbalanced datasets, such as the one used here, where the number of positive labels varies substantially from the number of negative labels, sensitivity (also known as recall), precision, and the F-score are often more informative metrics for insight into the model’s performance [21].

Evaluating the performance measures and the overall reliability of the prediction models in a clinical sense requires interpretation of the type of errors and associated cost of the errors in correctly identifying germline versus somatic variants. Assuming we label true germline mutations as “positive”, and true somatic mutations as “negative”, then the confusion matrix would result in the following errors: type 1 error (or false positive) which signifies the incorrect inference of a true somatic mutation as germline, and a type 2 error (or false negative), which signifies the incorrect inference of a true germline mutation as somatic. The cost of misclassifying a true somatic mutation (type 1 error), in a clinical sense, is equal to the cost of performing germline testing which can correct the incorrect inference results. However, the cost of failing to identify the presence of a germline mutation (type 2 error), may result in neglecting to validate the mutational status through germline testing and possibly leaving the treating physician without critical information that could alter the treatment strategy and missing the opportunity for cascade testing of at-risk family members. Therefore, two subsets of data were created to compute the performance measures. The first consisted of PGV along with pathogenic somatic variants, while the second consisted of all germline (PGV and VUS) as well as all somatic variants.

## RESULTS

The 1,467 eligible patients with both tumor sequencing results and independent germline sequencing (**Table 1, Figure 1**), were predominantly female (73%), were White/Caucasian non-Ashkenazi (84%) and had a median age of 54 years (range 1–88) at first primary tumor diagnosis. The most frequent tumor types were breast (22%), epithelial ovary including fallopian tube and peritoneum (21%) and colorectal cancers (19%). A total of 725 patients (49%) had reportable germline findings, 285 (29%) of whom had at least one PGV and 440 (61%) had one or more VUS exclusively. Individuals self-identified as Ashkenazi Jewish had a high rate of PGV (38 of 44; 13% of all PGV) in contrast with Hispanic individuals who had the lowest rate (0.7%) in our cohort. The approximate frequency of PGV in genes associated with the sequenced tumor were: small bowel carcinoma, 29%; urothelial carcinoma, 25%; renal cell carcinoma, 24%; colorectal carcinoma, 15%; breast carcinoma, 14%; epithelial ovarian carcinoma, 13%; and pancreatic adenocarcinoma, 13%. No PGVs were detected in 35 genes analyzed; 1–3 PGV were detected in 22 genes, and >3 PGV were detected in 13 genes (**Supplementary Tables 1, 2**).

**Figure 1:**
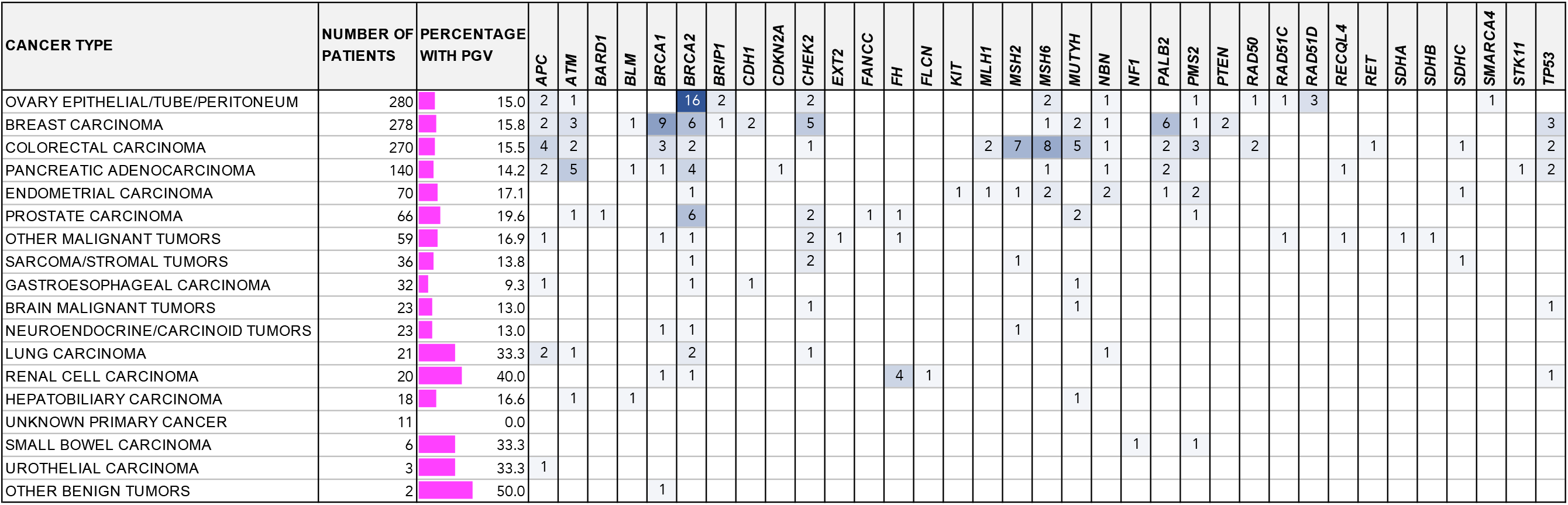
The parentage of tumors per cancer type with pathogenic germline variants in 35 genes from germline testing.

**Table 1.**
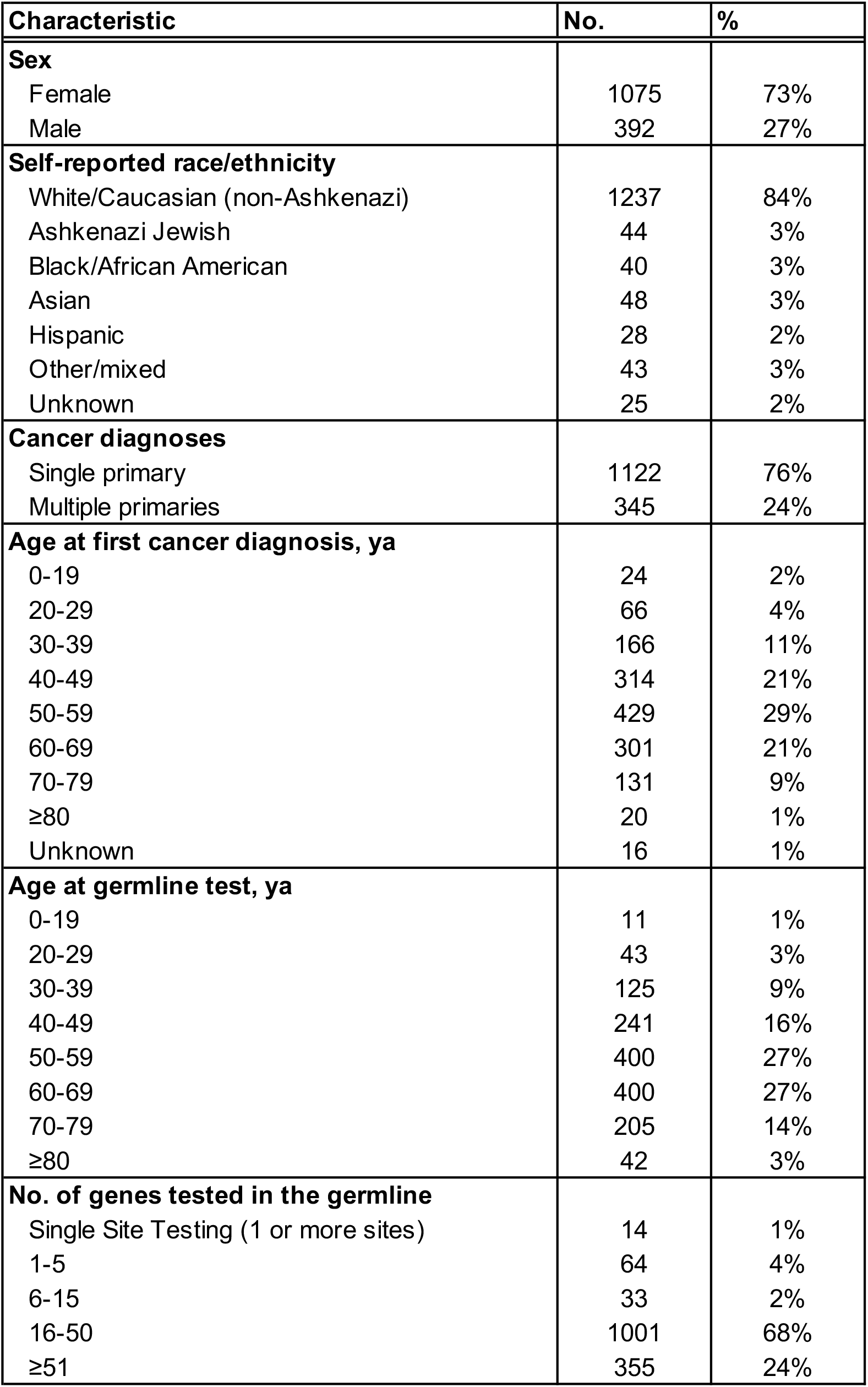
Patient Demographic and Clinical Characteristics (N = 1,467).

| Characteristic | No. | % |
| --- | --- | --- |
| **Sex** |  |  |
| Female | 1075 | 73% |
| Male | 392 | 27% |
| **Self-reported race/ethnicity** |  |  |
| White/Caucasian (non-Ashkenazi) | 1237 | 84% |
| Ashkenazi Jewish | 44 | 3% |
| Black/African American | 40 | 3% |
| Asian | 48 | 3% |
| Hispanic | 28 | 2% |
| Other/mixed | 43 | 3% |
| Unknown | 25 | 2% |
| **Cancer diagnoses** |  |  |
| Single primary | 1122 | 76% |
| Multiple primaries | 345 | 24% |
| **Age at first cancer diagnosis, ya** |  |  |
| 0-19 | 24 | 2% |
| 20-29 | 66 | 4% |
| 30-39 | 166 | 11% |
| 40-49 | 314 | 21% |
| 50-59 | 429 | 29% |
| 60-69 | 301 | 21% |
| 70-79 | 131 | 9% |
| ≥80 | 20 | 1% |
| Unknown | 16 | 1% |
| **Age at germline test, ya** |  |  |
| 0-19 | 11 | 1% |
| 20-29 | 43 | 3% |
| 30-39 | 125 | 9% |
| 40-49 | 241 | 16% |
| 50-59 | 400 | 27% |
| 60-69 | 400 | 27% |
| 70-79 | 205 | 14% |
| ≥80 | 42 | 3% |
| **No. of genes tested in the germline** |  |  |
| Single Site Testing (1 or more sites) | 14 | 1% |
| 1-5 | 64 | 4% |
| 6-15 | 33 | 2% |
| 16-50 | 1001 | 68% |
| ≥51 | 355 | 24% |

Tumor-only sequencing using the OncoPanel assay detected 5,426 variants across 70 cancer susceptibility genes; [22, 23]; matched germline testing results of the relevant gene was available for 3,988 of them. In total, 728 variants were detected by germline testing among which 231 were annotated as PGV and 497 as VUS. The remaining 3,260 variants were not reported in germline analysis and therefore were deemed to be somatic (**Figure 2A, Supplementary Table 1**); 1,792 of these variants (55%) were predicted to be likely pathogenic or pathogenic.

**Figure 2:**
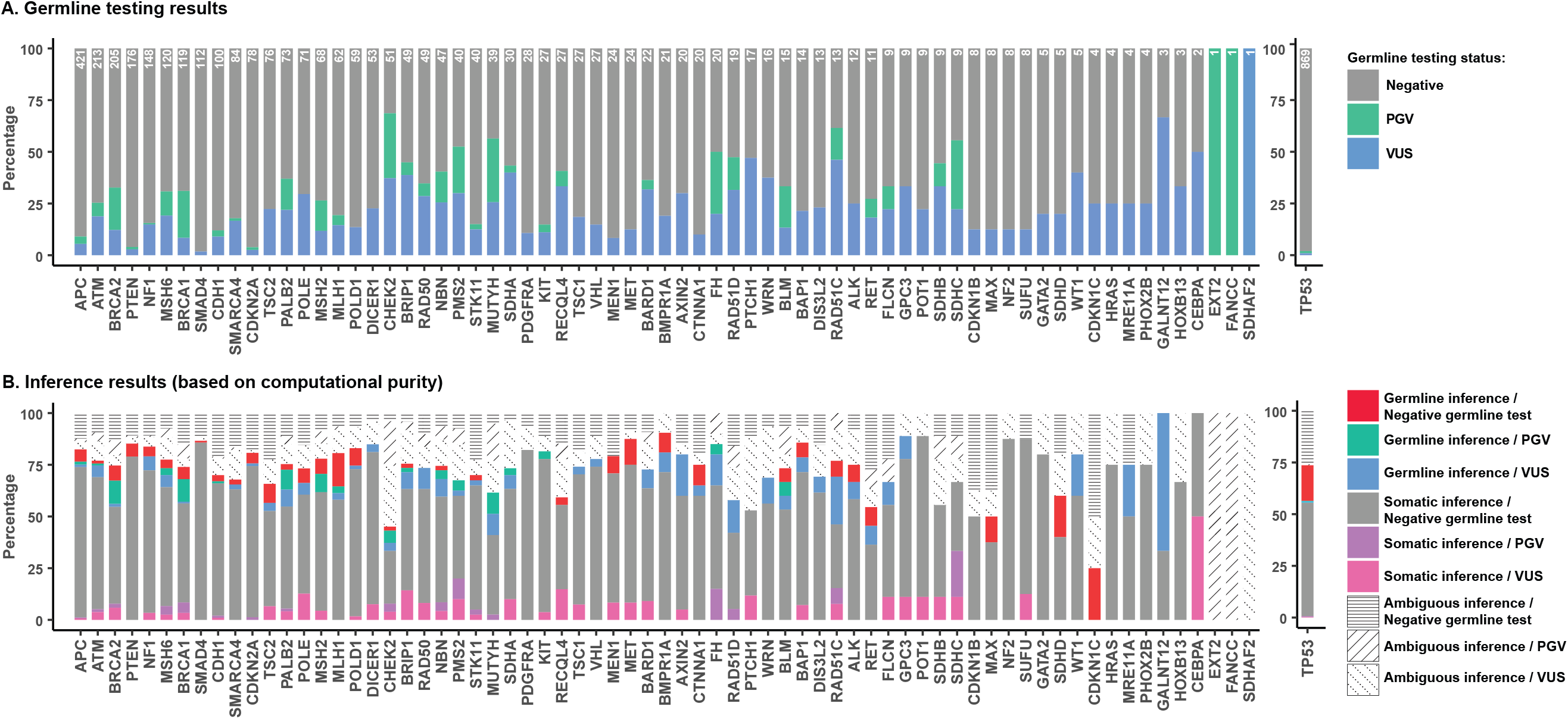
A) Matched germline testing results for 3,988 variants detected by tumor-only sequencing in 70 cancer susceptibility genes, including 231 PGV, 497 germline VUS, and 3,260 somatic variants. B) Inference of mutational status using computational purity estimates compared to germline testing results. Results using histological purity estimates are shown in **Supplementary Figure 2**.

We inferred non-ambiguous, germline or somatic mutational status for 3,028 (75.9%) variants using computational estimates of specimen tumor purity and 3,173 (79.5%) variants using histological estimates (**Figure 2B, Supplementary Figure 1**). Inferred mutational status using either purity estimate were highly concordant (Jaccard index = 0.84 [24]). We evaluated the accuracy of inference results considering all germline variants (PGV and VUS) or only the PGV, along with somatic variants. The performance results using computational and histological purity estimates were highly concordant (**Figure 3, Supplementary Figure 3**). For simplicity, the remainder of the results will only report those from computational estimates, which are calculated as a part of our pipeline.

**Figure 3:**
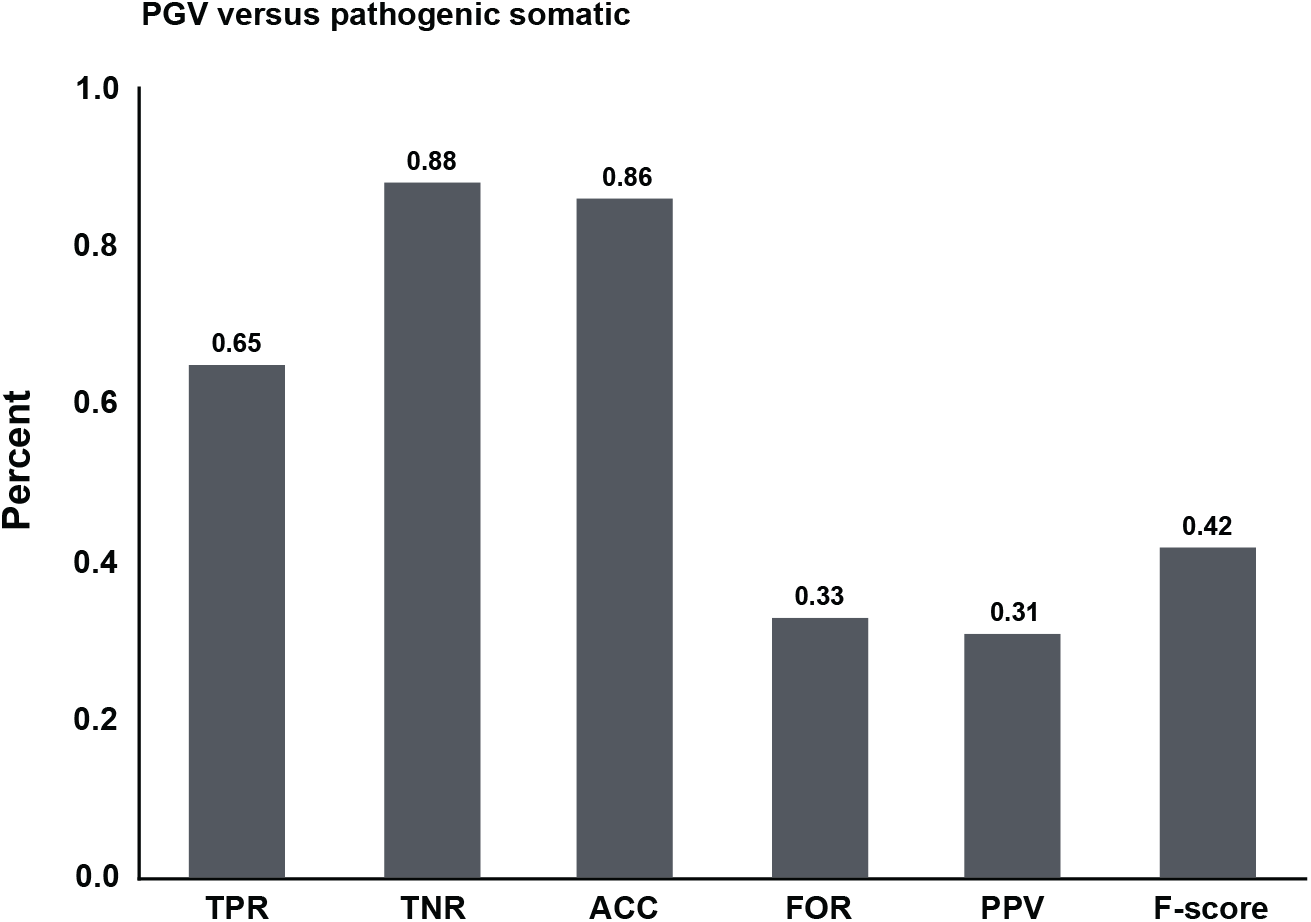
True positive rate (TPR or recall), true negative rate (TNR), accuracy (ACC), false omission rate (FOR), positive predictive value (PPV or precision) and F-score performance measures for the inferences made using computational purity for pathogenic germline variants (PGV) versus pathogenic somatic variants. Result from did not change when pathogenicity of variants was considered (**Supplementary Figure 3A**) or using histological purity estimates (**Supplementary Figures 3B, 3C**).

When only the PGV and pathogenic somatic variants were considered, the true positive rate (TPR or sensitivity) was 65%, signifying the rate at which the PGV were correctly inferred. The true negative rate (TNR or specificity), which indicates the rate of correctly inferring true somatic variants, was 88%. The false omission rate, indicating how often a true PGV was incorrectly inferred as somatic was only 3%. The positive predictive value (precision) and the F-score were 31% and 42%, respectively, which could be attributed to the relatively low number of true pathogenic germline variants in the dataset compared to the number of true somatic variants. The overall accuracy of the analysis was 86%. These results did not change when pathogenicity of variants was considered (**Supplementary Figure 3**).

The majority of somatic variants that were incorrectly inferred as germline (278 of 394, 71%) had VAF >50% (**Figure 4A**), while 83% of true pathogenic germline variants (118 of 143) that were incorrectly inferred to be somatic had VAF <50% (**Figure 4B**). In the latter group, 22% of incorrectly inferred variants corresponded to indels. There was no significant difference between the focal copy-number or the types of variants – SNV or indel – with correct or incorrect inference. The percentage of variants with ambiguous inference was 20.5% and 24.1% using computational and histological purity estimates, respectively. Variants with ambiguous inference had a mean VAF of 52.2% (**Figure 4C, Supplementary Figure 4**). Expected allele frequency for germline heterozygous mutations is 50% and is independent of tumor purity; however, various somatic models also predict expected VAF of 50% across a range of purity and copy-number values (**Supplementary Figure 1**), which could result in ambiguous inference.

**Figure 4:**
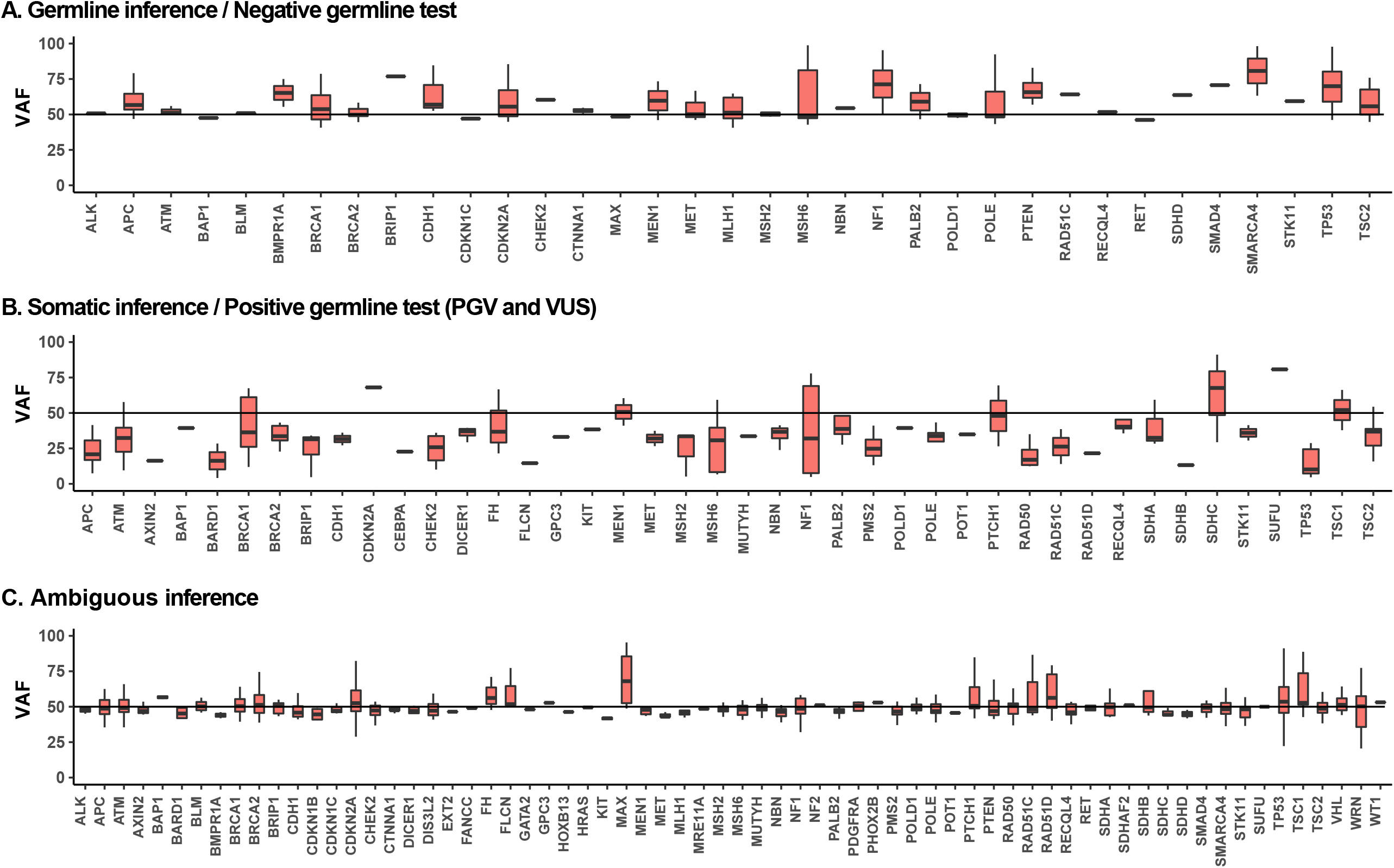
Allele frequencies distribution of variants with incorrect or ambiguous inference per gene: A) Somatic variants with germline inference. B) Germline variants with somatic inference. C) Germline and somatic variants without a statistical inference (ambiguous). Results using computational purity estimates are shown; results from histological purity estimates are shown in **Supplementary Figure 4**.

Mean sequencing depth of variants with correct predictions (mean = 295.5; standard deviation (sd) = 147.1) were not significantly different from those with incorrect inferences (mean = 283.7; sd = 149.6). Similarly, tumor purity estimates were not significantly different in specimen with variants that were inferred correctly (mean = 48.1, sd = 21.4) versus incorrectly (mean = 44.8, sd = 23.9). Although low sequencing depth and inaccurate purity estimation can contribute to the false inference of variants, they did not systematically bias the performance of our model.

Somatic mutations in *TP53* are the most common alterations in human cancers, whereas germline *TP53* mutations, the underlying cause of Li-Fraumeni syndrome (LFS), are rare. We correctly inferred mutational status of germline mutations in 5 of 5 LFS cases. Peripheral blood sequencing for germline testing was also positive for 3 additional cases; however, the VAFs of these variants in blood and tumor were 6–18%, suggesting detection of mosaicism due to clonal hematopoiesis [25]. Moreover, 17.6% (150 of 852) of *TP53* variants detected by tumor sequencing were falsely inferred to be germline. These variants were detected at VAFs significantly higher than their respective specimens’ estimated tumor purity (rank-sum test *p* <0.001, **Supplementary Figure 5**). Similarly significant patterns were also observed for incorrectly inferred somatic variants in *APC* (rank-sum test *p* = 0.018) and *PTEN* (rank-sum test *p* = 0.003), implying that inference of variants with high VAF in tumor suppressor genes may be affected by inaccuracies in estimating purity and confounded by unreported focal copy-number changes from loss of the wild-type allele or copy-neutral LOH.

Prior to inferring mutational status, the overall proportion of the PGV to all pathogenic variants detected by tumor-only sequencing was 11%, ranging from 1% to 100% for individual genes (**Supplementary Table 2**). When only the pathogenic variants with VAF >30% were considered [14], this ratio increased to 19%, resulting in a sensitivity (TPR) of 91% for detection of true germline variants, a specificity (TNR) of 50% for detection of pathogenic somatic variants, and an overall accuracy of 55%. In contrast, our model’s non-ambiguous, correct inference for 71% of pathogenic somatic variants increased the ratio of PGV to remaining pathogenic variants to 31%, without imposing any VAF criteria.

Next, we assessed the likelihood for the loss of the wild-type allele or copy-neutral LOH for all germline and somatic variants with correctly inferred mutational status. In total, a significantly larger percentage of PGV (72%) had LOH compared to 58% of germline VUS (chi-squared *p* <0.001) and 39% of pathogenic or likely pathogenic somatic variants (chi-squared *p* <0.001) (**Figure 5A**). The prevalence of PGV with LOH was evident when we focused on the genes associated with specific cancers, including both those with high and moderate/low penetrance.

**Figure 5:**
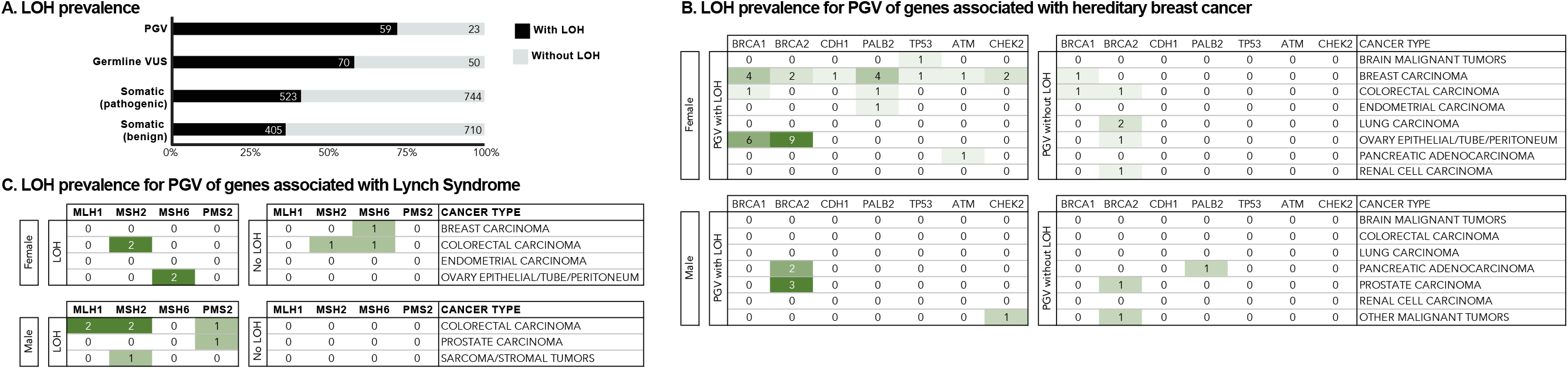
A) Percentage of PGV, germline VUS, and somatic pathogenic and benign variants with and without LOH in all genes. B) Prevalence of PGV with and without LOH per cancer type for genes associated with breast cancer in females and males. C) Prevalence of PGV with and without LOH per cancer type for genes associated with Lynch syndrome in females and males.

The high-penetrance genes associated with hereditary breast cancer include *BRCA1*, *BRCA2*, *CDH1*, *PALB2*, *PTEN*, *STK11*, and *TP53*, while *ATM* and *CHEK2* are considered as moderate/low-penetrance [26]. In females, LOH was demonstrated for *BRCA1* PGV and *BRCA2* PGV in 86% (6 of 7) of breast and 94% (15 of 16) of ovarian tumors, whereas LOH was demonstrated for only 33% of *BRCA1/2* PGV (2 of 6) in other tumor types (**Figure 5B**). LOH was demonstrated for all PGV (13 of 13) in *PALB2*, *TP53*, *ATM, CDH1* and *CHEK2* in all tumor types. In males, LOH was demonstrated for *BRCA2* PGV in 83% (5 of 6) of pancreatic and prostate tumors. These results agree with known prevalence of pathogenic germline alterations in these genes for corresponding cancer types [27-29].

The high-penetrance *MLH1*, *MSH2*, *MSH6* and *PMS2* genes are associated with Lynch Syndrome [30]. LOH was evident for 78% of the PGV (7 of 9) in colorectal cancers of both sexes. In females, LOH was demonstrated for both *MSH6* PGV in ovarian tumors (2 of 2) (**Figure 5C**). Overall, considering both males and females, 79% of PGV in the Lynch syndrome genes were found with LOH (11 of 14) across all tumor types. The results are again consistent with the status of pathogenic alterations associated with the Lynch syndrome particularly for ovarian cancer in females and colon cancer in males.

In contrast, somatic variants in the genes associated with the hereditary breast cancer or Lynch Syndrome did not show a significant correlation between inferred LOH and pathogenicity, although other events resulting in biallelic inactivation could not be ruled out.

## DISCUSSION

Estimates of the prevalence of inherited susceptibility to cancer are still imprecise in the general population. Emerging data from clinical sequencing assays indicate that the incidence of PGV may be as high as 17.5% in unselected cancer patients, and even higher for specific histologic types, including epithelial ovarian cancer and urothelial carcinoma [31, 32], reflecting the dependence on penetrance and tissue-specificity [33].

Extracting clinically relevant information from sequencing data requires accurate annotation of somatic and germline alterations by comparing a tumor’s molecular profile with the patient-matched normal samples. Paired tumor-normal analysis can also help identify somatic events that impact the genes with a PGV that may result in its biallelic inactivation [34]. However, most commercial and academic laboratories lack control germline DNA analysis and produce reports that may not address whether or not a variant is actually somatic. In this study, we presented a gene-independent bioinformatics workflow that, using commonly available measurements from tumor sequencing (i.e. total depth, focal ploidy, and VAF) can select the most likely germline versus somatic mutational status and assess evidence for loss of heterozygosity. By analyzing each variant in the context of specimen purity, we eliminate the need for *ad hoc* VAF criteria [14], or complex analyses of raw sequencing data [35]. We validated our approach using available germline testing results from 1,608 cancer patients. Where pathogenic variants across 70 cancer susceptibility genes were detected in tumor sequencing, inference of pathogenic germline variants had an overall sensitivity of 65%, specificity of 89%, and accuracy of 85% using computational purity estimates with highly concordant results from histological estimates.

Our performance statistics established a balance between the ability to detect the germline mutations (sensitivity) and the somatic mutations (specificity). Gene-specific, VAF-based criteria for identifying patients with PGV from tumor-only data could be highly sensitive; however, their application also results in a high number of type 1 errors, and thus, low specificity and overall accuracy [14]. In our data, only 11% of all detected pathogenic variants were PGV. In contrast, accurate inference of status for 71% of true pathogenic somatic variants led to a three-fold increase in the proportion of PGV to remaining pathogenic variants.

Following the Knudson two-hit model, tumorigenesis in PGV carriers is caused by the presence of a heterozygous germline alteration followed by the somatic loss of the remaining wild-type allele in the tumor cells by genomic alterations, or more rarely epigenetic silencing [36]. As not all cancers that arise in carriers may be driven by the germline alteration, it is important to determine whether a germline variant is accompanied by loss of the wildtype allele in a given cancer, both to understand the pathogenesis and to guide therapy. Our results showed a significant association between pathogenicity of germline alterations and the loss of the wild-type allele, highlighting the importance of distinguishing biallelic and LOH events from monoallelic PGV as a biomarker for therapeutic response [37]. In particular, the high rates of inferred LOH for pathogenic *BRCA1* and *BRCA2* variants in breast and ovarian cancers in our data are consistent with similar findings using other sequencing platforms suggesting existence of selective pressures for biallelic inactivation in these tumors [29, 34].

Systematically, the lower the sequencing depths at which a particular variant is detected, the lower the confidence in accuracy of measuring its VAF. Clinical tumor-only sequencing assays are mandated to have a relatively high depth of sequencing compared to research-grade whole-genome and whole-exome platforms; therefore, they are capable of identifying SNV and indels with high confidence. Sequencing at depth of coverage >300x is expected to provide sufficient power to accurately measure allele abundances and to statistically assess potential germline origin and zygosity of detected variants [18, 38]. In fact, with an average coverage depth of ~290x in our data, we did not observe a systematic difference in sequencing depth or specimen purity between the variants with true or false inferences. While germline variants with incorrectly inferred somatic status had VAF <50%, somatic variants with incorrectly inferred germline status had VAF >50%, highlighting the dependency of our approach on accurate VAF measurements. This lower than expected VAFs of PGV in tumor-only sequencing data suggests either a problem in variant calling, undetected low-level amplification of the wild-type allele or possibly presence of reversion mutations [19, 39, 40]. The high VAF of the confirmed somatic variants that were inferred to be germline suggests an over-estimation of tumor purity, computationally and histologically, in these samples. Although VAF for indels may be confounded by misalignment and variant calling inaccuracy, they were not associated with correct or incorrect inference, highlighting the utility of our approach for all variants with a measured VAF. Finally, our user-friendly, interactive bioinformatics application is freely available for academic use for performing these analyses on sequencing results from assays routinely employed in the clinic.

## CONCLUSION

While concurrent tumor and germline sequencing analyses for all cancer patients may become the standard of care in the future, the need to have an objective and reliable means of selecting patients for clinical germline testing confirmation is needed in clinical practice today. The increasing use of tumor-only sequencing assays can identify mutations in known cancer predisposition genes, raising the possibility of germline mutations and potentially the need for independent germline DNA assessment. Our analysis demonstrates that patients with potentially pathogenic germline alterations in cancer predisposition genes can be identified by analyzing their tumor-only sequencing data, suggesting that when a PGV is detected in the tumor specimen, the patient should be considered for genetic counseling and germline testing. Computational inference of LOH status for both germline and somatic variants may also be helpful in defining tumor pathogenesis and guiding therapy of individual cancers, even in the setting of paired germline and tumor sequencing data.

---

**Supplementary Figure 1:**
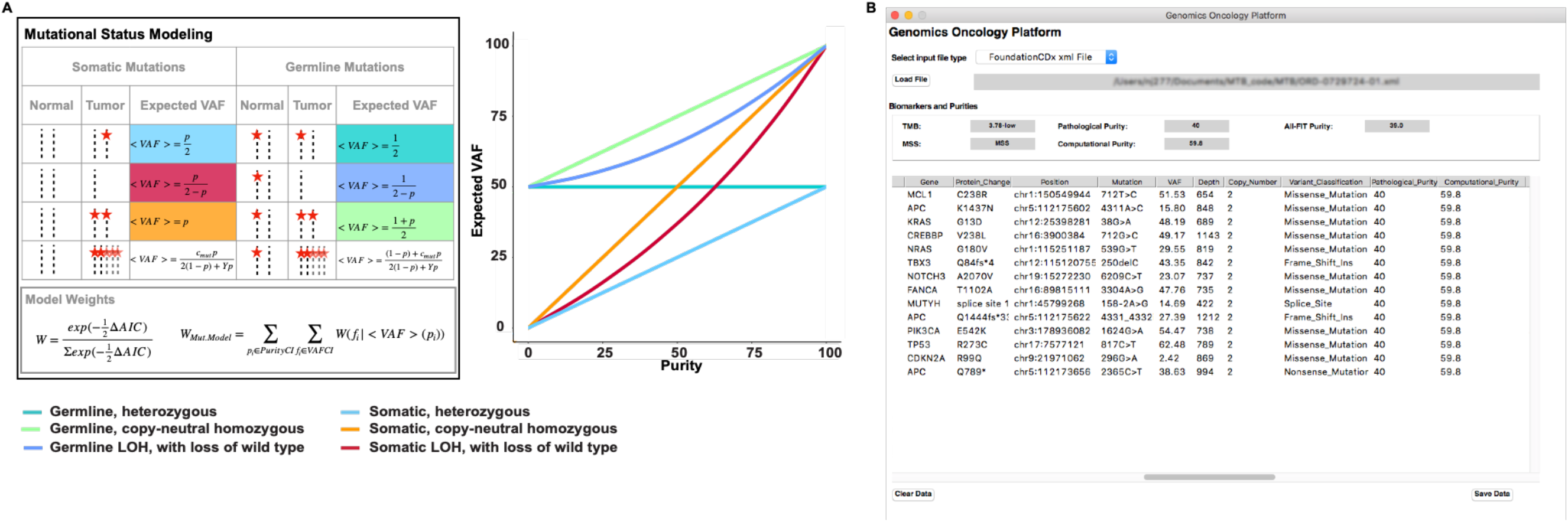
A) Expected variant allele frequencies as a function of purity for different mutational models (adapted from Khiabanian et al. JCO PO 2020). Akaike Information Criterion (AIC) weights are used to compare the likelihood of somatic and germline mutational models using the observed VAF and copy-number (ploidy) and sequencing depths at their positions. B) A snapshot of the application, showing variant data and status.

**Supplementary Figure 2:**
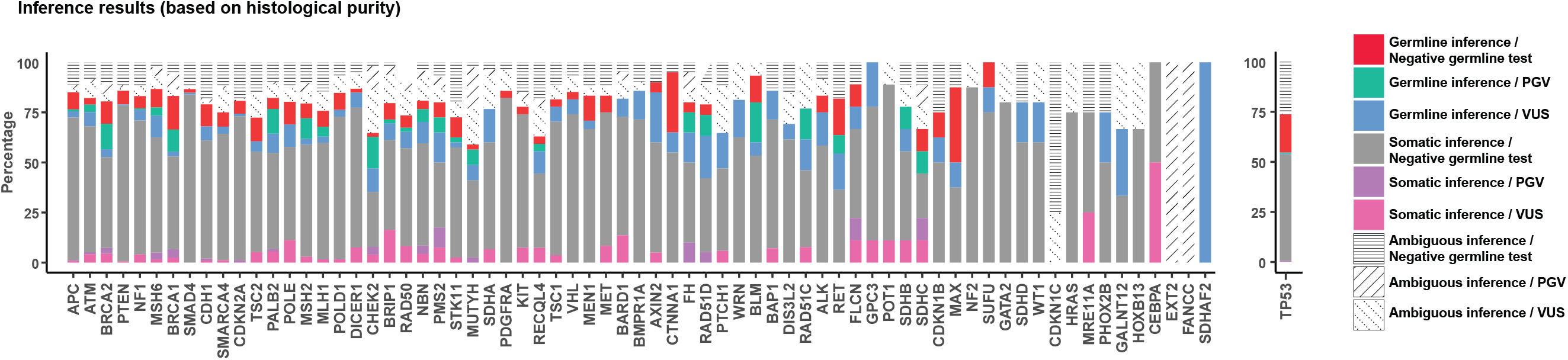
Inference of mutational status using histological purity estimates compared to germline testing results.

**Supplementary Figure 3:**
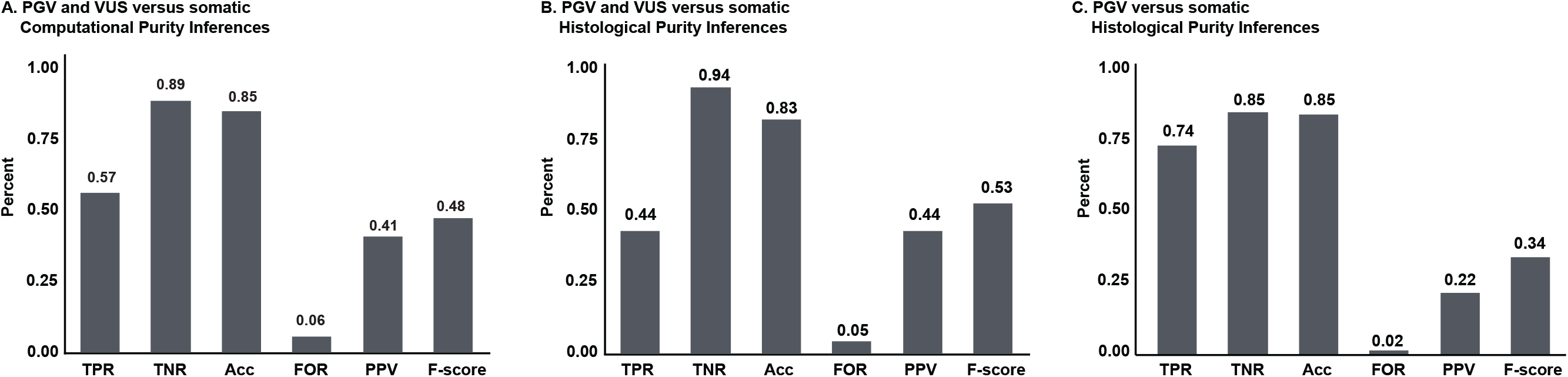
A) True positive rate (TPR or recall), true negative rate (TNR), accuracy (ACC), false omission rate (FOR), positive predictive value (PPV or precision) and F-score performance measures for the inferences made using computational purity for all germline variants (PGV and VUS) versus all somatic variants. B) Overall performance measures using histological purity for all germline variants (PGV and VUS) versus all somatic variants. C) Overall performance measures using histological purity when pathogenicity was considered for germline and somatic variants.

**Supplementary Figure 4:**
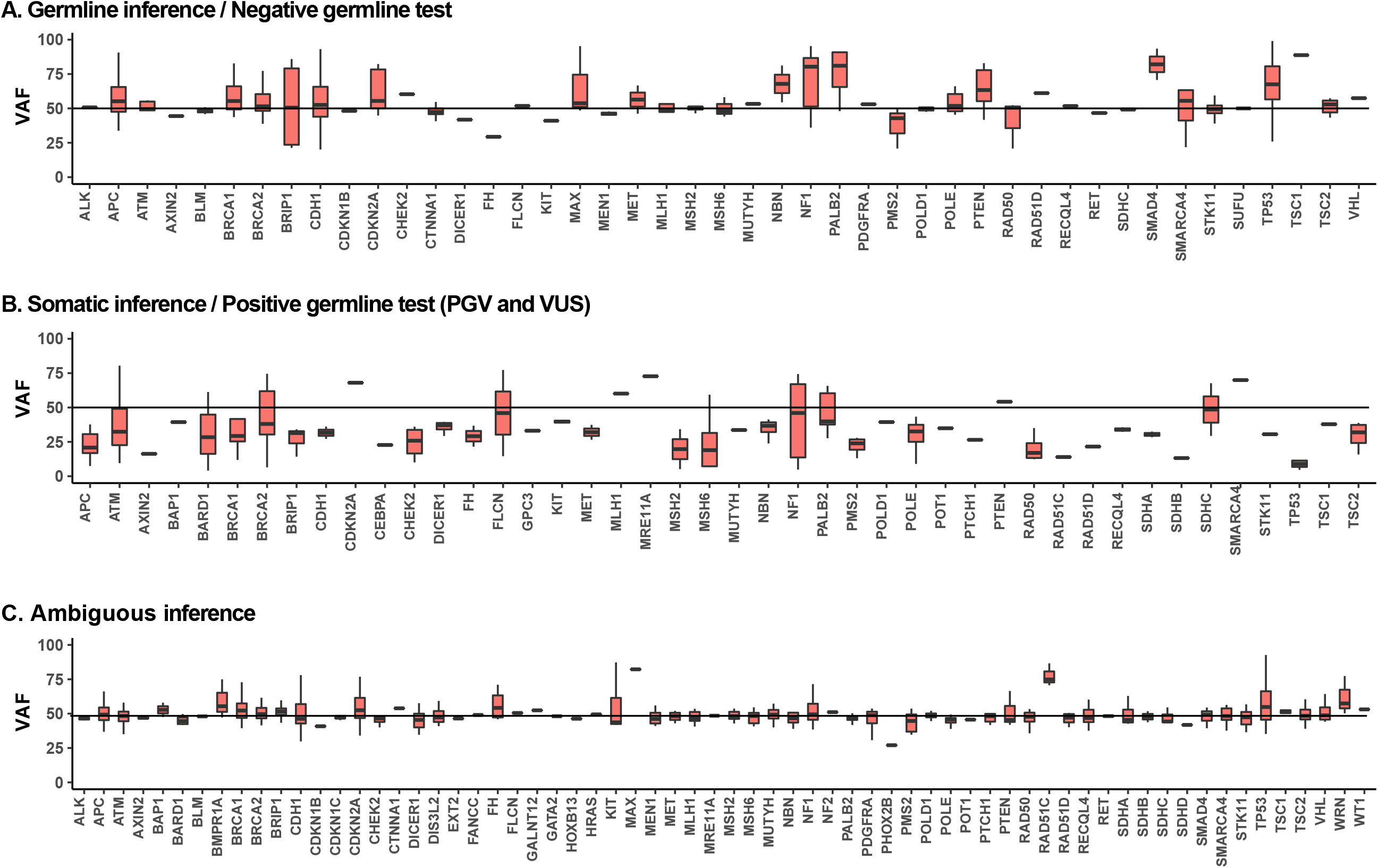
Allele frequencies distribution of variants with incorrect or ambiguous inference per gene: A) Somatic variants with germline inference. B) Germline variants with somatic inference. C) Germline and somatic variants without a statistical inference (ambiguous). Results using histological purity are shown.

**Supplementary Figure 5:**
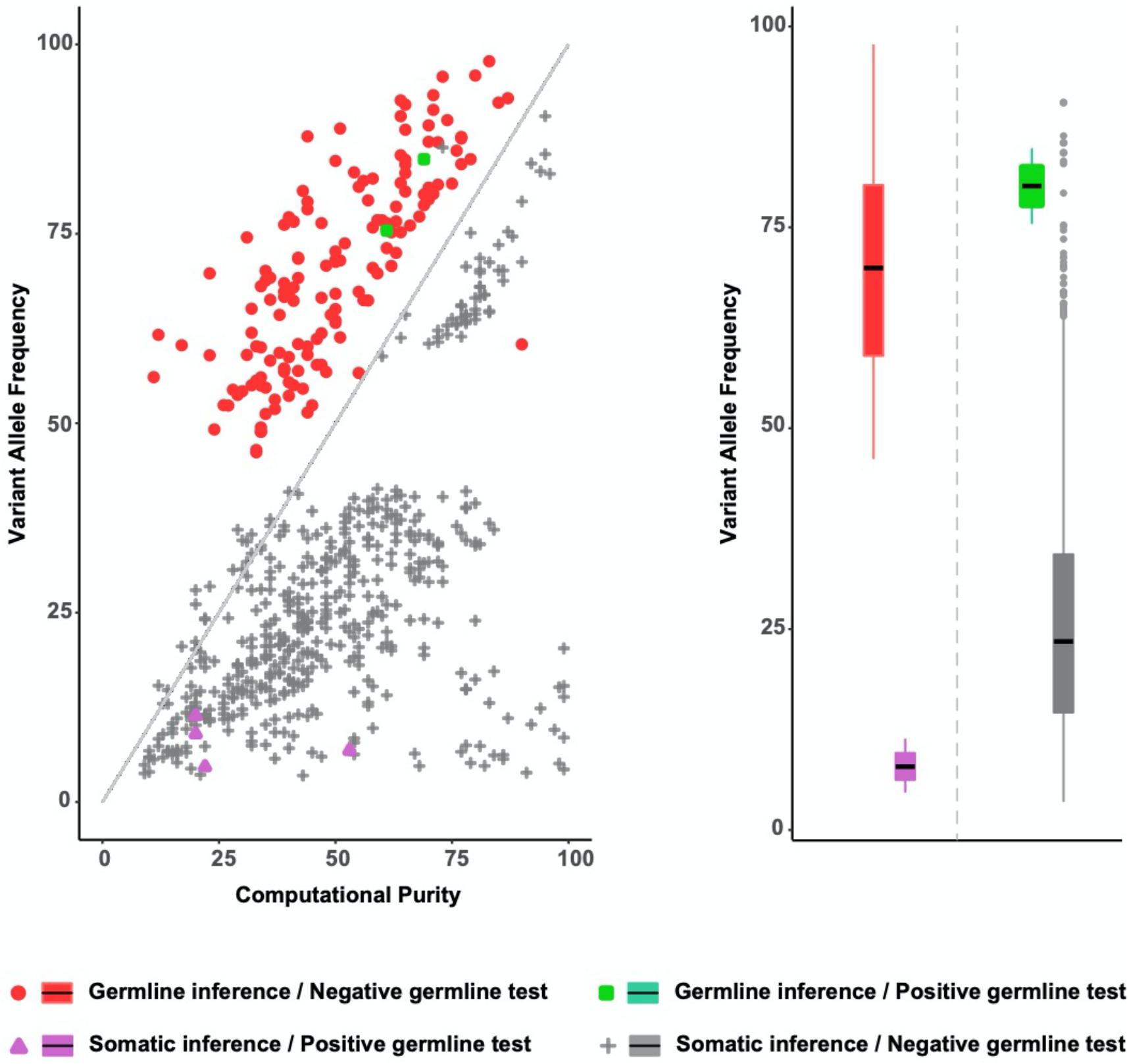
Variant allele frequency of *TP53* variants versus computational purity estimates, grouped based on correct and incorrect inferences.

